# Somatic evolution of stem cell mutations in long-lived plants

**DOI:** 10.1101/2025.01.13.632685

**Authors:** Frank Johannes

## Abstract

Long-lived perennial plants accumulate numerous somatic mutations with age. Mutations originating in stem cells at the shoot apex often become fixed in large sectors of the plant body due to cell lineage drift during repeated branching. Understanding the somatic evolution of such mutations requires knowledge of the effective stem cell population size, the cellular bottleneck strength during branch initiation, and the mutation rate. Here we show that these parameters can be estimated directly from cell-layer-enriched DNA sequencing data, thus filling a gap where no other *in vivo* method exists.

---

Many perennial plants exhibit extraordinarily long lifespans, ranging from centuries to millennia. Trees are a good example of this. Even at an advanced age they grow vigorously, iteratively forming new branches that produce leaves and fruits. All new aerial plant structures emerge from the shoot apical meristem (SAM), a population of cells situated at the tip of each growing shoot (Lyndon (1998)). The SAM consists of stem cells and other meristematic cells, which continually divide, either to self-renew or to differentiate. Somatic mutations that arise during stem cell divisions are particularly consequential as they can be propagated through the plant architecture over time, creating substantial intra-organism genetic heterogeneity. There is considerable interest in understanding the adaptive significance of this heterogeneity, as well as the mechanisms that contribute to the rate and patterns of mutation accumulation within the plant (Schoen and Schultz (2019)).

In higher plants, new lateral branches are initiated from a small number of precursor cells at the periphery of the SAM (Burian et al. (2016), Shi et al. (2016)). Their cell lineages trace back to a group of “apical stem cells” (ASCs) at the SAM center. The ASCs are distinct from surrounding cells due to their spatial positioning and slow mitotic division rate, and thus constitute an important unit of analysis (Burian (2021)). The limited number of precursors involved in branching creates cellular bottlenecks, allowing mutations in ASCs to become fixed in large sectors of the plant structure. This process has been termed somatic genetic drift (Reusch et al. (2021)).

Such fixed somatic mutations sometimes confer advantageous phenotypic effects on specific plant parts or modules, a phenomenon that has been extensively used in horticulture and fruit crop breeding, where mutant branches with desirable traits are propagated through techniques such as grafting, cuttings, or tissue culture (Foster and Aranzana (2018)). Advantageous somatic mutations have also been observed in natural settings (e.g. Padovan et al. (2013)), though they are likely rare. By contrast, deleterious ASC mutations are much more common (Eyre-Walker and Keightley (2007)), and contribute to a gradual build-up of mutational load over time (Scofield and Schultz (2006)). The harmful effects of these mutations are often hidden, however, because they typically arise as recessive alleles in a heterozygous state (Schoen and Schultz (2019)). As a result, the somatic evolution of ASC mutations is thought to be largely shaped by neutral somatic drift during branching (Klekowski and Kazarinova-Fukshansky (1984b), Chen et al. (2024), Yu et al. (2020), Tomimoto and Satake (2023)), rather than by selection acting on differences in fitness among cell lineages or plant modules (Folse and Roughgarden (2012), Antolin and Strobeck (1985), Klekowski and Kazarinova-Fukshansky (1984a), Pineda-Krch and Fagerstrom (1999), Otto and Orive (1995)).

Nonetheless, deleterious somatic mutations can be subject to strong selection once they enter the haploid gametic phase of the plant life cycle, where their effects are fully exposed. Mutations that escape purging during this stage can be transmitted to offspring, and so contribute to inbreeding depression at the population level (Lesaffre (2021)). Long-lived plant species appear to mitigate this problem by maintaining high outcrossing rates and large population sizes (Scofield and Schultz (2006)), both of which reduce inbreeding coefficients. Interestingly, an additional strategy utilized by long-lived plants to limit the accumulation of somatic mutations is to reduce the mutation rate itself (Lanfear et al. (2013), Johannes (2024), Shahryary et al. (2020)). This rate reduction seems to be particularly pronounced in ASC-derived cell lineages that eventually form the gametes (Goel et al. (2024)).

Recent DNA sequencing studies, mainly in tree species, have shown that the yearly rate of fixed somatic mutations is at least an order of magnitude lower than that observed in annual plants (Johannes (2024)). This trend is also evident in the lower evolutionary substitution rates detected in long-lived angiosperms and gymnosperms compared to short-lived species (De La Torre et al. (2017)). It has been proposed that these reduced rates are achieved by limiting the number of mitotic divisions in ASCs per unit time (e.g. Petit and Hampe (2006), Lanfear et al. (2013), Burian (2021), Johannes (2024)), thereby decreasing the likelihood of errors during DNA replication. An alternative, yet unexplored, hypothesis is that long-lived species have larger effective stem cell populations or recruit a greater number of precursor cells during branch initiation, both of which would dilute the effects of somatic drift and reduce the rate of fixation. Theoretical models predict that larger stem cell populations could also buffer the impact of deleterious mutations, should they display some level of dominance (Klekowski and Kazarinova-Fukshansky (1984a)).

To begin to test these hypotheses and to study the somatic evolution of ASC mutations from a quantitative perspective, detailed knowledge of the effective stem cell population size, the cellular bottleneck during branch initiation, and the mutation rate are needed for diverse plant systems. Despite recent technological advances in live imaging (e.g. Bradamante (2024)) and Crisper-based cell lineage tracing (e.g. Lu et al. (2024)), direct measurements of these parameters remain experimentally challenging. A major obstacle is to obtain measurements *in vivo*, and in a manner that is applicable across a wide range of plant species, including those that cannot be easily transformed genetically. Moreover, neither technology provides information about mutation rates.

One promising alternative strategy is to develop theoretical models of shoot branching that describe how these parameters affect changes in the frequency of ASC mutations in the SAM cell population over time. As these mutation frequencies can be measured directly using bulk sequencing of selected somatic tissues, fitting the model to such data could yield statistical estimates of the unknown parameters. Although theoretical models of somatic evolution have been developed, none are currently designed to be applied to actual somatic sequencing data for parameter estimation. This is particularly true of early theoretical models that focused on mutant cells as units of analysis (e.g. Klekowski and Kazarinova-Fukshansky (1984b), Klekowski and Kazarinova-Fukshansky (1984a), Otto and Orive (1995), Pineda-Krch and Fagerstrom (1999)). More recent models do have a DNA focus, but their application appears to require single-cell data of individual SAMs (Iwasa et al. (2023), Chen et al. (2024)), which is technically infeasible at the moment, or they make strong numerical assumptions about the very parameters that require estimation (Tomimoto and Satake (2023), Yu et al. (2024)).

Here we overcome many of these challenges and demonstrate that a modeling approach in combination with bulk sequencing of somatic cells is feasible. We begin by laying out the biological framework underlying the theoretical branching model. We then develop the model step-by-step and show how it can be used to derive and compute the expected frequency spectrum of ASC mutations in the SAM cell population. Finally, we provide a proof-of-principle demonstration of how parameter estimates can be obtained by fitting the model to recent layer-enriched DNA sequencing data of an apricot tree.

## Biological framework

The shoot apical meristem (SAM) is a zone of actively dividing cells at the shoot tip, driving the formation of new stems, leaves, and flowers. In seed plants, the SAM is organized into a tunica-corpus structure, with the tunica comprising the outer cell layers and the corpus forming the inner region. Most species possess a tunica that consists of one or two layers (L1, L2) where cells divide anticlinally (perpendicular to the surface). The underlying corpus (L3) has cells dividing in multiple directions (Lyndon (1998)) (Fig. S1). Stem cells in the SAM have the capacity for self-renewal and for producing progenitor cells that differentiate (Heidstra and Sabatini (2014)). Live imaging and cell lineage tracing in Arabidopsis and tomato (Burian et al. (2016)) have identified 3-4 resident stem cells at the very tip of the SAM surface that distinguish themselves from surounding cells by their extremely slow division rates (Lyndon (1998), Kwiatkowska (2008))(Fig.1A). These so-called apical stem cells (ASCs) are the origin of all SAM cell lineages, including those forming lateral organs and the germline (Burian (2021)). A similarly arranged, but independent, population of ASCs is present in the L2 and the inner L3 layer, just below the SAM surface. We will only consider the L1 case further, but similar arguments holds for the other layers.

### The cellular basis of branching

In higher plants, lateral branches arise from axillary buds, which form in the axils of developing leaves along the stem (Fig. S2). These buds originate from clusters of cells called axillary meristems (AMs) (Fig.1A, Fig. S2). AMs develop from a small group of precursor cells that are sequestered from the flanks of the SAM (Burian et al. (2016), Shi et al. (2016)), a process known as the “detached meristem model” of shoot branching (Wang and Jiao (2018)). These precursor cells retain meristematic properties within the leaf axil, gradually proliferating to form active AMs that later produce new lateral branches (Fig. S2). Once established, these AMs function as the SAM of the emerging shoot, possessing the same structural organization, self-maintenance capabilities, and ability to generate new organs as the original SAM (Nicolas and Laufs (2022)). In this way, shoot branching reflects a developmental shift, where the SAM of the primary shoot gives rise to new SAMs on lateral branches (Fig.1B, Fig. S2).

**Figure 1:**
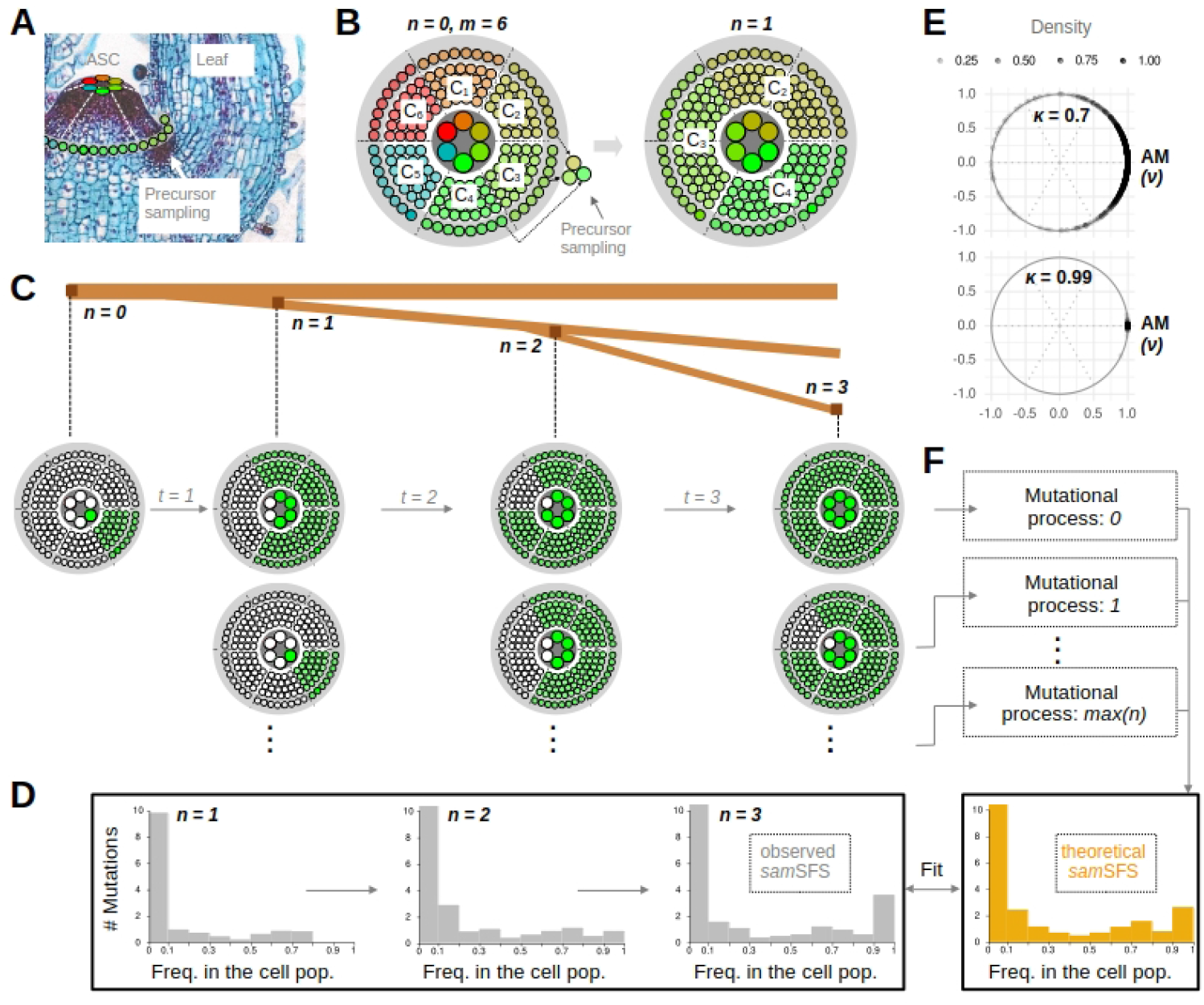
Biological basis and conceptual framework. **A**. Side-view of the shoot apical meristem (SAM). The colored circles at the tip represent six (*m* = 6) apical stem cells (ASC). Clonal descendants of these ASCs are shown in corresponding colors at the SAM periphery. The arrow points to the site of axillary meristem (AM) establishment. **B**. A schematic top-view of the SAM. Six clonal sectors (*C*_1_, …, *C*_6_) are shown at *n* = 0, with each sector corresponding to one of the ASC at the tip. At first branching (*n* = 1), precursors are “sampled” from *C*_2_, *C*_3_ and *C*_4_ during AM establishment, and give rise to a new SAM. The cellular composition of the new SAM reflects the proportional sampling of these sectors. **C**. Cell lineage composition of the SAM across three nested branching events. Here, lineages from clonal sector *C*_4_ become fixed in the SAM over time. In time intervals *t*_1_, *t*_2_ and *t*_3_ the ASC self-renews. Mutations in ASCs that occur in these intervals follow the fate of the cell lineages in which they arise. There is a new mutational processes, starting with each new branching iteration *n*, giving rise to mutations that are nested within those that appeared earlier. **D**. The frequencies of these mutations within the SAM cell population defines the site frequency spectrum (*sam*SFS) at *n*. **E**. Modeling of precursor sampling using a wrapped density distribution. The distribution governs the location of AM establishment (*ν*) and the density spread of selected precursor around the site of AM establishment (*κ*). Sampling can be highly localized (bottom) or more spread around the site of AM establishment (top). **F**. Modeling of precursor sampling along with the mutational processes during nested branching yields the theoretical *sam*SFS, which can be fitted to the observed *sam*SFS for parameter estimation.

### Somatic drift and cell lineage fixation

In the time interval between two branching events, the ASC population self-renews through asymmetrical cell divisions (Fig. S3). Each apical stem cell produces two descendant cells, where one retains its stem cell function at the central position of the SAM, while the other is displaced toward the SAM periphery, where it undergoes a series of symmetrical cell divisions (Fig. S3). In the L1 layer, this latter process generates spatially defined clonal sectors in the SAM, with each sector having its origin in one of the apical stem cells at the center (Burian et al. (2016)) (Fig.1A,B). The sampling of L1-derived precursor cells at the SAM periphery is therefore an indirect sampling of ASCs, albeit via their clonal descendants. Depending on the number of clonal sectors and the degree of spatial constraints in AM establishment, precursor sampling can be a source of substantial somatic drift (Reusch et al. (2021), Chen et al. (2024)), leading to ASC lineages becoming either fixed or lost in the SAM over time (Fig.1C). In the limiting case, AMs are initiated from a single cell from each of the SAM layers. In this scenario, there is immediate fixation of one clonal lineage in the new SAM. However, if precursor sampling is more “relaxed”, involving cellular contributions from multiple clonal sectors, fixation may require several nested branching iterations to reach fixation (Fig.1C).

### Mutation accumulation in the SAM

The developmental fate of *de novo* mutations within the SAM is closely related to the fate of the cell lineages in which they arise. We focus exclusively on mutations within the ASC population. We assume that these mutations occur with some rate *µ* (per bp per year) during the self-renweal phase between branching events, and that their frequencies in the SAM evolve neutrally through somatic drift. To see this, consider a mutation that arises in an apical stem cell in the first internode time interval *t*_1_ (Fig.1C). This mutation essentially “barcodes” all cell lineages that derive from this cell, and its fixation dynamics is therefore equivalent to the fixation of the cell lineage that carries it.

However, the interval *t*_1_ is not the only interval contributing mutations to the SAM. New and independent mutations also arise in ASCs in any subsequent branching intervals *t*_2_, *t*_3_, … *t*_*n*_, generating mutational “barcodes” that are nested in those acquired in earlier intervals (Fig.1C). The continuous flow of *de novo* mutations into the SAM, coupled with somatic drift, therefore generates a distribution of mutational frequencies in the SAM during branching, with developmentally older mutations being already fixed at high frequency in the SAM population, and more recent mutations appearing at intermediate to low frequencies. We define the complete mutation frequency distribution as the SAM site frequency spectrum (*sam*SFS) (Fig.1D). The shape of the *sam*SFS at any point in time holds statistical information about the underlying branching process, including the mutation rate (*µ*), the size of the ASC population (*m*) as well as the spatial constraints during precursor sampling (*κ*).

In what follows we develop a theoretical model that incorporates these parameters (Fig.1E,F). Using the theory we derive the theoretical *sam*SFS, and show that it can be fitted to layer-enriched DNA sequencing data for parameter estimation.

## Theoretical model and parameter inference

### Unit circle representation of the SAM

Let us represent the L1 surface layer of the SAM as a unit circle in polar coordinates (Fig.1E). The unit circle is defined as:

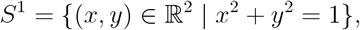

which describes all points (*x, y*) on the boundary of a circle with a radius of 1 in the two-dimensional plane. We assume that the unit circle is divided into *m* cell partitions, labeled *C*_1_, *C*_2_, …, *C*_*m*_ (Table 1). Each partition corresponds to a subpopulation of clonal cells descended from a single ASC at the center of the

**Table 1:** Definition of important model notation.

| Symbol | Definition |
| --- | --- |
| $m$ | Number of apical stem cells (ASCs) in the SAM |
| $n$ | Branching iteration index ($n = 0, 1, 2, \dots$) |
| $C_i^{(n)}$ | Clonal sector $i$ in the SAM at branching iteration $n$. Cells in this clonal sector are all descended from ASC $i$. |
| $\phi_i^{(n)}$ | Proportion of SAM cells at iteration $n$ descended from ASC $i$; directly measures the fraction of a SAM layer occupied by $C_i^{(n)}$ |
| $t_n$ | Internode time interval ending at branching iteration $n$, measured in years or in number of ASC cell divisions ($n = 0, 1, 2, \dots$) |
| $\nu$ | Mean direction (location) of AM initiation along the SAM periphery |
| $\kappa$ | Concentration parameter of the wrapped distribution; governs spatial constraint during precursor selection and thus indirectly the cellular bottleneck strength during branching |
| $\mathcal{F}_i$ | Fixation time: branching iteration when $\phi_i^{(n)} \approx 1$ |
| $\mu$ | Somatic mutation rate (per base, per year) |

SAM (Fig.1B). The angular width of each partition *i*, denoted by Δ*θ*_*i*_, is defined as the difference between its starting angle *a*_*i*_ and its ending angle *b*_*i*_:

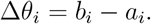

We suppose that at *n* = 0, when the SAM is first established during development, all *m* partitions are of equal size (Fig.1B). Hence, the initial angular width for each sector is:

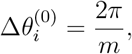

with the starting angle 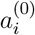 and the ending angle 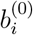 given by:

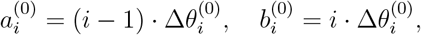

where *i* = 1, …, *m*. The uniform partitioning of clonal sectors at *n* = 0, implies that the initial proportion, 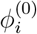 (Table 1), of each sector 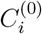 is simply:

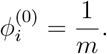

### Branching: precursor sampling and re-normalization

Branching is initiated by the formation of AMs at the SAM periphery (Fig.1A,B). It starts with the selection (i.e. sampling) of a small number of peripheral cells to act as precursors. For simplicity, several previous models assumed that this sampling is uniform (Klekowski and Kazarinova-Fukshansky (1984b), Klekowski and Kazarinova-Fukshansky (1984a), Klekowski et al. (1985), Otto and Orive (1995), Chen et al. (2024), Yu et al. (2024), Pineda-Krch and Fagerstrom (1999), Antolin and Strobeck (1985)), so that all peripheral cells have an equal chance of being selected. In reality, the sampling processes is spatially constrained, often involving only cells that are in close proximity of each other. The degree of this constraint can depend on factors such as the precise SAM topology and the actual number of precursors, all of which can vary across species (Schnablová et al. (2017)). To account for these spatial aspects Tomimoto and Satake (2023) proposed to model precursor sampling using a wrapped distribution on a unit circle (Fig.1E). A wrapped distribution is derived by taking a standard linear distribution and “wrapping” it around the circle to ensure that the random radian variable *θ* remains confined to the interval [0, 2*π*). The general form of the probability density function (PDF) of a wrapped distribution is:

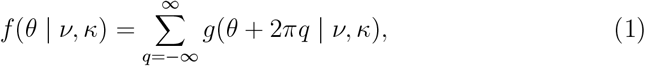

where *g*(· |*ν, κ*) is the probability density function of a linear distribution on the real line. This base distribution defines how probability mass is allocated before wrapping and determines the overall shape and concentration of the resulting circular distribution. Here, *q* is an integer index that represents how many times the real line has been “wrapped” around the circle Kato and Jones (2013). As can be seen, the distribution depends on two parameters, *ν* and *κ* (Table 1). Both have important biological interpretations. The parameter *ν* is the location parameter and gives the “mean direction”; that is, the angle around which the distribution is centered. Biologically, this sets the location of AM initiation and lateral bud formation along the SAM periphery (Fig.1E). The parameter *κ* is the “concentration” parameter. It controls how tightly the probability mass is concentrated around *ν*. In biological terms, *κ* determines the degree of spatial constraint in precursor selection. For small *κ*, the distribution is spread out more uniformly around the circle, while for large *κ*, the distribution becomes tightly concentrated around *ν* (Fig.1E). Indirectly, the parameter *κ* thus quantifies the cellular bottleneck during precursor sampling.

Now, consider the first branching event at *n* = 1. We are interested in the probability that cells from clonal partitions 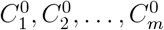 are selected as precursors for AM initiation at some location *ν*^(1)^ along the SAM periphery. This probability can be derived for each *C*_*i*_ by integrating Equation 1 over the angular range of each partition:

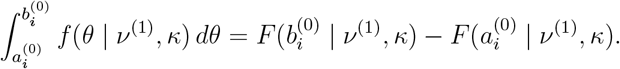

where *F* (· | *ν, κ*) is the cummulative distribution function (CDF), 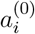 and 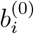 are the angular start and end points of the *i*-th partition just prior to branching, respectively, and *ν*^(1)^ is the location of AM initiation. As the AM matures into a bud and forms a new SAM on the emerging branch, the clonal partitions have to be re-normalized to account for the new cellular proportions:

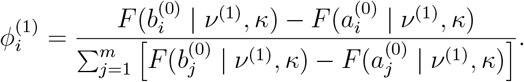

where the summation in the denominator ensures that the total probability across all 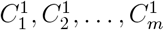 remains 1. These updated proportions thus present the proportion of SAM cells following branching at *n* = 1, whose lineages trace back to apical stem cell *i* at *n* = 0.

### Nested branching and cell lineage evolution

The above arguments suggest the following recurrence relation for iterative nested branching events:

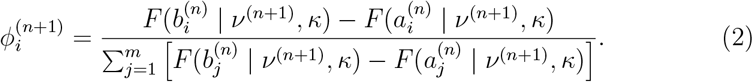

with updated boundaries:

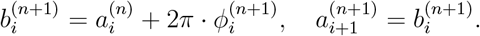

Notice that we allow the mean direction *ν*^(*n*)^ of the wrapped distribution to change with each branching event *n*. This should account for the fact that the location of AM initation along the SAM periphery is not fixed over time. We model the dynamic behavior of *ν*^(*n*)^ in two ways. First, we assume that *ν*^(*n*)^ rotates around the unit circle according to the following deterministic function:

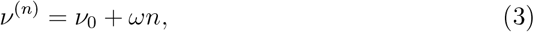

where *ω* is a fixed angular velocity. This model can capture different forms of phyllotaxy; that is, the periodic placement of leaf primordia and their accompanying AMs along the axis of shoot growth (Kuhlemeier (2017)). For example, we can have *ω* = *π* so that AM initiation is shifted by a half a circle with each branching iteration, thus completing a full rotation around the SAM with every second branching event. A second model assumes that *ν*^(*n*)^ is randomly chosen, at each branching iteration, from a uniform distribution on the interval [0, 2*π*), i.e.,

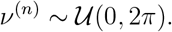

This model is realistic if one considers that not all AMs actually develop into lateral branches. It could therefore reflect some level of stochasticity in AM activation or dormancy. In addition, the spatial coordinates of the clonal sectors in the newly established SAM are not fixed, but re-established randomly with respect to the site of AM initiation.

The evolution and convergence of the recurrence Equation 2 can be evaluated numerically using:

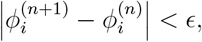

where *ϵ* is a small positive number. Note, that this also implies that the SAM partition boundaries stabilize:

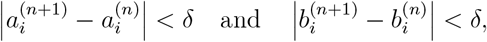

where *δ* is a small positive number.

### Fixation and fixation times

We seek to explore the conditions, at convergence, under which the proportion *ϕ*_*i*_ for a given partition reaches a state of effective “fixation”, meaning it approaches or effectively reaches a value close to 1. For a given partition 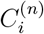, we define fixation as:

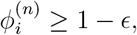

where *ϵ* is a small positive number, indicating that 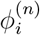 is “sufficiently close” to 1 to be considered fixed. Now, let ℱ_*i*_ denote the fixation time for partition *C*_*i*_ (Table 1). This is defined as the smallest branching iteration *n* for which the fixation criterion is first met:

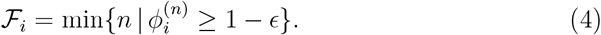

While the above derivation focuses on the fixation of cell lineages that originate in ASC *i*, there is a direct connection with the fixation of *de novo* mutations. As already argued above, a mutation that first arises in ASC *i* in time interval *t*_1_ “barcodes” all cell lineages that derive from this ASC, and its fixation dynamics is therefore equivalent to that of the cell lineages themselves. Since the lineages of only one of the *m* ASCs converges to fixation, the probability of a mutation in a random ASC going to fixation is therefore simply *m*^−1^ (Klekowski and Kazarinova-Fukshansky (1984b)).

### Modelling the *sam*SFS

We can build on the preceding analysis to derive a theoretical form of the site frequency spectrum (*sam*SFS) of somatic mutations that originate in apical stem cells (ASCs). The *sam*SFS characterizes how these mutations are distributed across frequencies within the shoot apical meristem at a given branching iteration *n*. Crucially, this frequency distribution is shaped by the nested structure of meristem development and mutation accumulation. To formalize this, we generalize the recurrence framework introduced earlier and allow for multiple, temporally distinct mutation-drift processes to evolve simultaneously. Each process *k* = 1, …, *n* begins at a different branching iteration and corresponds to mutations arising in interval *t*_*k*_ (Table 1). These processes share a common wrapped sampling distribution at each iteration but are initialized independently. For each process *k*, the unit circle representing the SAM is initially divided into *m* equal clonal sectors (see (Fig.1C)):

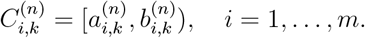

The initial proportions of each 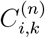 are uniform: 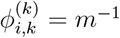. At each subsequent branching iteration, the wrapped distribution *f* (*θ*; *ν*^(*n*)^, *κ*) - with mean direction *ν*^(*n*)^ and concentration parameter *κ* - governs the evolution of these proportions, so that the update rule becomes:

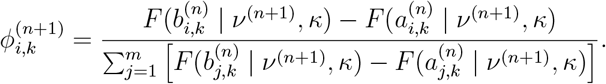

The output of each process *k* is a vector proportions of the form:

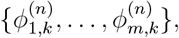

which represent the allele frequencies of mutations arising in each of the *m* ASCs during *t*_*k*_. To construct the *sam*SFS, we partition the interval (0, 1] into *u* non-overlapping bins:

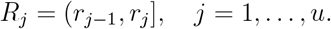

The frequency of SAM cells descended from apical stem cell *i* under mutation process *k*, is assigned to one of these bins using:

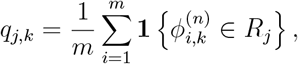

where **1** {·} denotes an indicator function. To reflect the expected number of mutations contributed by each branching interval, we scale the contribution of each component by its expected number of mutations. This expectation is given by:

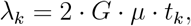

where *G* is the size of the callable genome, *µ* is the somatic mutation rate (per base per year), and *t*_*k*_ is the duration of interval *k* (Table1). These expectations are normalized to form the mixture weights:

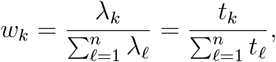

under the assumption that *G* and *µ* are constant across intervals. This formulation ensures that longer time intervals contribute proportionally more mutations to the site frequency spectrum than recent ones, all else being equal. Hence, the resulting *sam*SFS is a finite mixture distribution over bins, with mixture weights *w*_*k*_ and component-specific probabilities *q*_*j,k*_. The marginal probability that a mutation falls into bin *R*_*j*_ is:

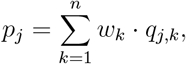

defining a probability vector **p** = (*p*_1_, …, *p*_*u*_) used in subsequent inference.

### Model fitting: single sample analysis

The observed data consist of *N*_mut_ somatic mutations, each with a measured variant allele frequency (VAF). These VAFs are usually re-scaled (e.g., multiplied by 2 in diploid systems) to obtain a measure of the proportion of cells that carry a given mutation. We discretize the re-scaled VAFs into the *u* bins *R*_1_, …, *R*_*u*_, resulting in a count vector:

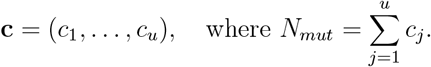

Given the theoretical mixture model defined by **p**(Ψ), where Ψ = (*m, κ, µ*) denotes the model parameters, we assume:

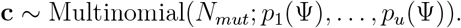

The corresponding likelihood of the somatic mutation data is:

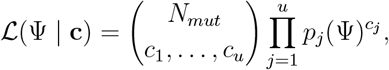

and the log-likelihood (up to a constant) becomes:

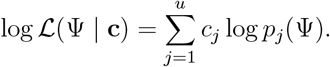

To estimate the parameters, we maximize the log-likelihood:

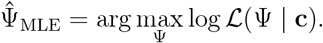

### Joint Sample Analysis

When multiple samples are available, we consider two strategies for joint inference. The first strategy performs likelihood-based integration across samples by maximizing a weighted average of the sample-specific log-likelihoods. This approach accounts for sample-specific branching histories but treats samples independently given shared parameters. Let *S* be the number of samples, then we can write

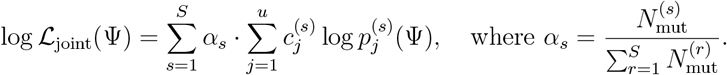

The maximum likelihood estimate is then obtained by:

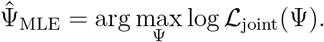

The second strategy aggregates information across samples by combining mutation counts and internode durations (*t*_1_, …, *t*_*k*_) for samples that share the same number of branching intervals *k*. This yields a single composite dataset in which the mutation signal is enhanced through pooling. For each interval *t*_*k*_, let 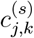 denote the number of mutations in bin *R*_*j*_ from sample *s*. The aggregated mutation counts are defined as:

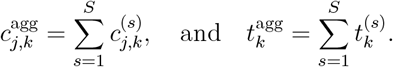

We compute the aggregated mixture weights as:

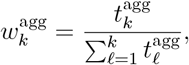

and the normalized bin frequencies within each interval:

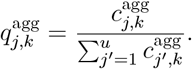

The resulting aggregated *sam*SFS is:

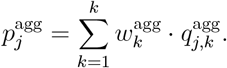

This formulation is particularly useful in low-mutation-rate regimes, where mutation counts within individual samples are sparse and statistical power can be gained by combining across time and samples. Inference in this setting mirrors the single-sample analysis, with the distinction that the dataset represents a temporally and mutationally aggregated sample. Parameter estimation proceeds by maximizing the log-likelihood for the aggregated count vector 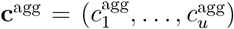, defined analogously:

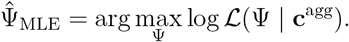

While both joint analysis strategies provide mechanisms to integrate information across samples, care must be taken to ensure that the samples are sufficiently independent in their developmental histories. This consideration is particularly important for the aggregation-based approach, where mutation counts and branch durations are pooled under the assumption of independence. If samples share extensive ancestral branch paths (e.g., neighboring branches with a recent common branch point), it may introduce systematic biases (Johannes (2025)). To avoid this bias, aggregation should be restricted to samples with largely disjoint developmental lineages, or alternatively, shared ancestry must be explicitly modeled when present.

### Incorporating Mutation Burden via Penalized Likelihood

In all models described above, the mutation rate parameter *µ* is absent from the normalized site frequency spectrum, due to its cancellation when computing the mixture weights. As a result, inference based solely on the frequency distribution does not constrain *µ*, rendering it non-identifiable from the multinomial likelihood alone. To enable joint estimation of *µ* alongside structural parameters such as the number of apical stem cells *m* and the concentration parameter *κ*, we introduce a penalty term based on the expected mutation burden under the model. Let *N*_mut_ denote the observed number of somatic mutations across all bins, and let 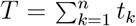 be the total developmental time across all branching intervals. The expected number of mutations under parameter *µ* is then:

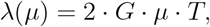

where *G* is the size of the callable genome. We define an *L*_2_ penalty that measures the squared deviation between the observed and expected mutation burden:

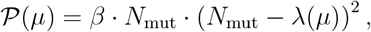

where *β* is a user-defined scaling constant that governs the strength of the penalty (default: *β* = 1). The resulting penalized log-likelihood used in inference becomes:

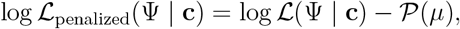

where Ψ = (*m, κ, µ*) and log ℒ (Ψ | **c**) denotes the standard log-likelihood associated with any of the single-sample, joint-sample, or aggregated models. This formulation imposes a biologically informed constraint on *µ*, anchoring its estimate to the total number of observed mutations. Importantly, accurate estimates of *κ* and *m* can be obtained even if *β* = 0 or if *N*_mut_ is biased by a constant factor (e.g., due to consistent false positive or false negative rates in variant ascertainment), as long as the shape of the *sam*SFS is unaffected.

## Results and discussion

### Fixation times of apical stem cell mutations

We computed the expected fixation time of ℱ a *de novo* mutation occuring in an apical stem cell, given that the clonal lineage of that cell had gone to fixation. Note that “fixation time” is defined here as in Equation 4 to mean the number of branching events, rather than absolute time. Our goal was to assess to which extent ℱ varies as a function of the ASC population size (*m*) and the concentration parameter (*κ*). To perform these computations, we assumed a wrapped Cauchy distribution underlying Equation 2 (see Methods). We explored a param- eter regime, where *κ* varied over a grid from 0.7 to 0.999 for two different values of *m* (4 and 40). The lower bound grid value of *κ* (0.7) was chosen such that at least 90% of density of the wrapped Cauchy was contained within the interval [0, *π*] of the unit circle (see Methods). In other words, precursor sampling was restricted to one half of the SAM to reflect the fact that AM formation is not dispersed around the SAM periphery. Finally, we considered two models for AM initiation (Fig.2A). The first model sampled the location of AM initiation along the SAM periphery from a uniform distribution at each branching iteration, while the second model sampled according to a phylotaxic process in steps of 0.5 · *π* (see Theory section).

**Figure 2:**
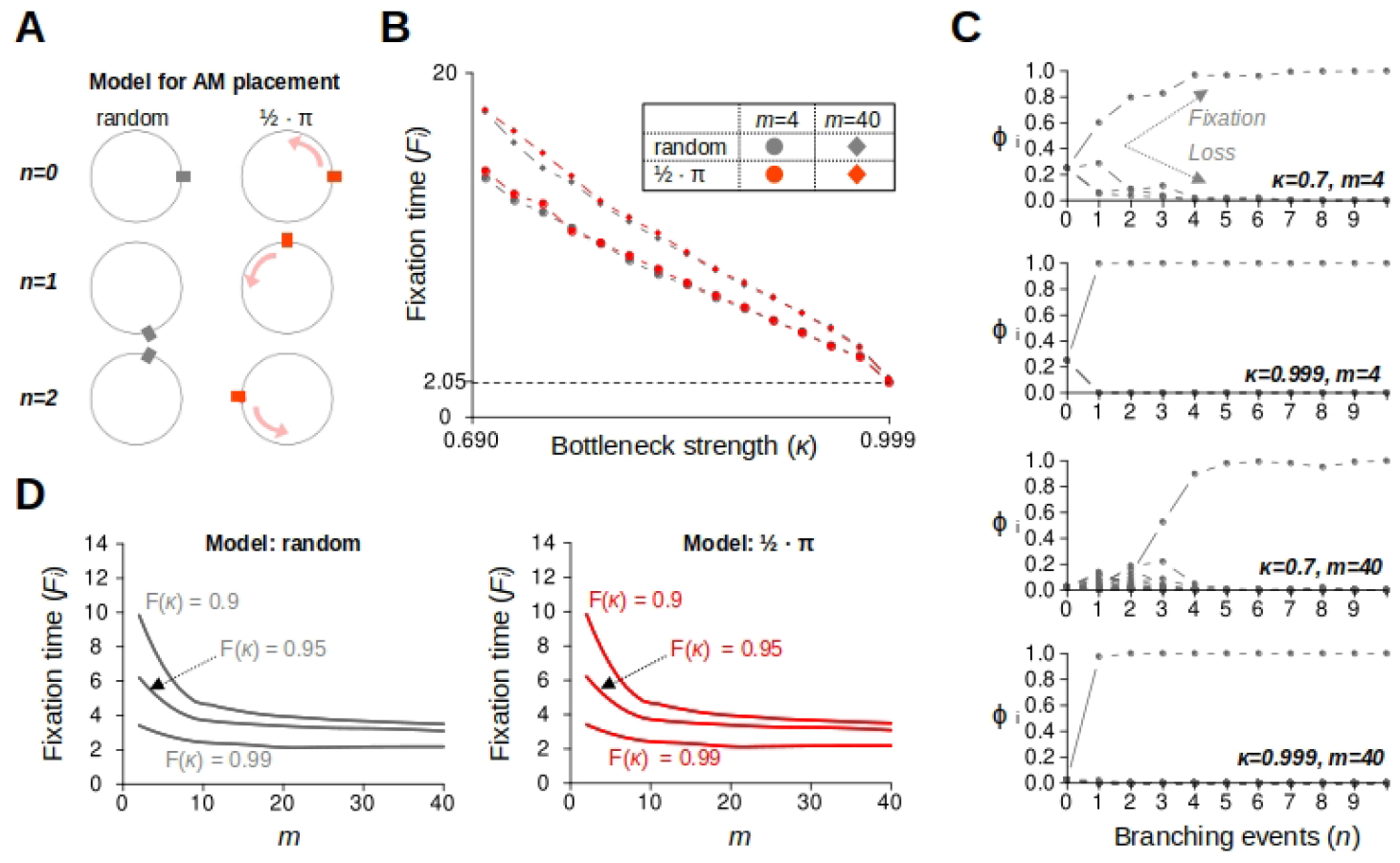
Fixation times of apical stem cell mutations. **A**. Two models of axillary meristem (AM) placement. In the first model, AMs are established at random locations along the SAM periphery with each branching iteration. In the second model, AM locations are sampled according to a phylotaxic process in steps of 0.5 · *π*. **B**. Computation of fixation times as a function of the concentration parameter *κ* and the size of the ASC population *m* for the two AM placement models. Fixation times denote the number of nested branching events that are required for a cell lineage that originates in one of the ASC cells at *n* = 0 to reach a relative frequency of 1 within the SAM. These fixation times also hold for *de novo* mutations that have arisen in the ASC self-renewal phase *t*_1_. **C**. Shown are several numerical realizations of the stochastic processes that leads to the fixation (and loss) of cell lineages. The proportion *ϕ*_*i*_ (y-axis) of SAM cells that are descendants of apical stem cell *i* at *n* = 0, plotted as a function of the number of nested branching events (x-axis). Fixation is relatively slow in situations where *κ* is relaxed (*κ* = 0.7) and *m* is large (*m* = 40), leading to intermediate frequency in early branching iterations. **D**. Fixation times become nearly independent of *m* as *κ* → 1. The reason is that precursor sampling becomes progressively restricted to a single clonal sector, creating strong somatic drift. In the limiting case, only a single precursor cell is chosen for AM establishment.

We found that fixation times are mainly influenced by the concentration parameter *κ*. For *κ* = 0.7, fixation was reached after approximately 12 to 18 nested branching events, but after only *≈* 2 for *κ* = 0.999 (Fig.2B,C). Without providing formal proof, it is intuitive that when *κ* → 1, the sampling convergences to a single-cell bottleneck, so that ℱ → 1, as only one clonal sector will be sampled. Interestingly, the impact of *m* on the fixation time was relatively modest. Its largest effect occurred for small values *κ* and became gradually less prominent with increasing *κ*. This observation points at an interesting relationship between *m* and *κ*: the impact of ASC population size *m* becomes negligible as soon as the concentration parameter is large enough to ensure that precursor sampling is restricted to only one of the *m* SAM clonal sectors. To illustrate this point, we assessed different values of *m* ranging from 2 to 40, and calculated the corresponding *κ* such that 90%, 95% and 99% of the total Cauchy density was contained within the angular width, Δ*θ* = 2*π* · *m*^−1^, for each *m* (see Methods). As expected, we found that fixation time ℱ becomes nearly independent of *m* as *κ* → 1 (Fig.2D). Finally, a comparison between the random and the phyllotaxy models did not reveal notable differences for any of the parameter combinations, although we detected slight fixation delays when phyllotaxy was accounted for (Fig.2B,D, Table S1).

### *sam*SFS dynamics during repeated nested branching

Recent work in *A. thaliana* suggested that branching is initiated from only one precursor cell of each of the L1, L2 and L3 layers (He et al. (2024)). However, it is currently unknown if this holds generally across species. Our analysis shows that if spatial sampling is more relaxed, the somatic fixation of ASC mutations would require several branching iterations (e.g. Fig.2C). As new mutations arise continuously during ASC self-renewal there will always be some mutations that originate in the internode interval just prior to the latest branching event, or in the one just before that and so on. The result is a mixture of mutation frequencies within the SAM, ranging from older mutations that are already fixed (frequency of 1) to more recent mutations that are present in only a few cells (frequency close to zero) (Chen et al. (2024)). As described in the Theory section, we refer to the complete frequency distribution as the *sam*SFS.

It is intuitive that the *sam*SFS is shaped by rate of *de novo* mutations (*µ*) as well as somatic drift (via *m* and *κ*). As the number of nested branching iterations (*n*) becomes large (*n*→ ∞), the *sam*SFS should converge to a mutation-drift equilibrium. However, this limiting case never occurs in reality, because plants initiate relatively few nested branching events during their life-time. In tree species, for instance, the life-time number of nested branching events is usually no more than five or six. Moreover, recent DNA sequencing studies searching for somatic mutations in trees, particularity in fruit trees, have sampled terminal branches that had experienced as few as two nested branching events (e.g. Goel et al. (2024)). It is therefore important to understand the temporal dynamics of the *sam*SFS in finite time; that is, over time-scales that are empirically relevant.

To begin to motivate how the *sam*SFS changes during iterative nested branching, we considered a hypothetical branch that had undergone 1, 2, 3, …, 10 equally spaced nested branching events. We assumed that the total age of the branch is 10 years, so that the internode age is e.g. five years when only two branching events occurred, and one year when 10 branching events. This ensured that the total number of *de novo* mutations that arise over time is independent of the number of branching events. Similar to the previous section, we considered two distinct values for the concentration parameter *κ* (0.7 and 0.99) to reflect relaxed and strong spatial constraint in precursor sampling, respectively, as well two very different values for the size of the ASC population *m* (4 and 40).

We found that, in the case where the spatial constraint is relatively relaxed (*κ* = 0.7) and the ASC population large (*m* = 40), a considerable proportion of somatic variants appear at low to intermediate frequencies in the SAM population, particularly in early branching iterations (Fig.3A). However, the shape of the *sam*SFS changes progressively with increased branching iterations, with low to intermediate frequency variants either becoming fixed or lost due to drift (Fig.3A). A similar, but less pronounced pattern, can be seen when the ASC population is small (*m* = 4). As expected, the situation is different for large *κ* (e.g. *κ* = 0.99). In this case, somatic drift efficiently clears low to intermediate variants, even after only a few nested branching events, and generates a characteristic U-shaped SFS (Fig.3B). That said, low frequency variants do persist in this case, especially when the ASC population is large *m* = 40 and the number of branching events is small.

**Figure 3:**
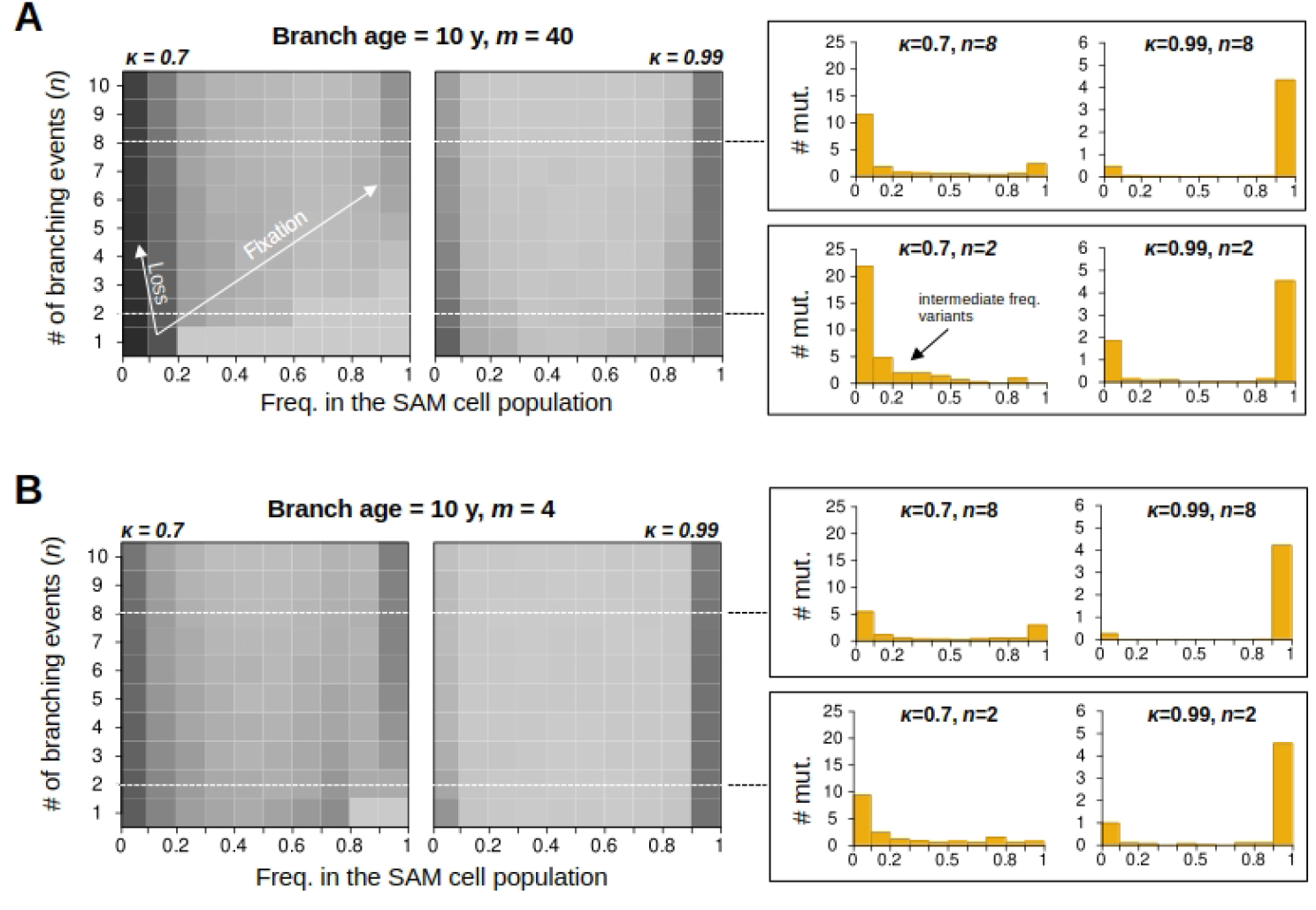
*sam*SFS dynamics during nested branching. Computation of the *sam*SFS assuming a 10 year old branch, a haploid mutation rate of 10^−9^ per bp per year, and a haploid genome size of 240 Mb. **A**. Given a relatively large ASC population size (*m* = 40), shown are the frequency changes of the *sam*SFS during successive nested branching events under a relaxed and strong drift regime (*κ* = 0.7 vs. *κ* = 0.99, respectively). Each row is a 1D projected histogram (see right panels), with dark gray indicating high- and light gray indicating low density. **B**. As in A.), but assuming a relatively small ASC population size (*m* = 4). Right panels: For each scenario, the *sam*SFS is highlighted after two (*n* = 2) and after eight branching events (*n* = 8). While drift rapidly generates a U-shaped *sam*SFS in the course of repeated nested branching, mutations do appear at intermediate frequencies in the SAM cell population. This phenomenon is most pronounced in early branching iterations (e.g. *n* = 2), when drift is weak (*κ* = 0.7) and the ASC population is large (*m* = 40).

The above insights reflect recent observations made by genomic studies in plants, which found a wide-range of low and high frequency variants (e.g. Schmitt et al. (2024)), even after layer-enriched sequencing was taken into account (e.g. Goel et al. (2024), Amundson et al. (2023)). The utility of the *sam*SFS is that it can be applied to such sequencing data to obtain parameter estimates of *m, κ* and *µ*, given that *n* and the ages of the internode segments is known.

## Data application: Simulation study

To be able to apply our method to empirical data, we implemented the likelihood inference framework described above (section: Modelling the *sam*SFS) in a software package (https://github.com/jlab-code/samSFS). We first evaluate the performance of the likelihood-based inference framework by conducting simulations in which synthetic datasets were generated under known model parameters and then reanalyzed using the full inference pipeline (Fig. 4). The goal was to assess how accurately the method could recover the true values of the number of apical stem cells (*m*), the spatial concentration parameter (*κ*), and the mutation rate (*µ*) across a range of biologically plausible conditions.

**Figure 4:**
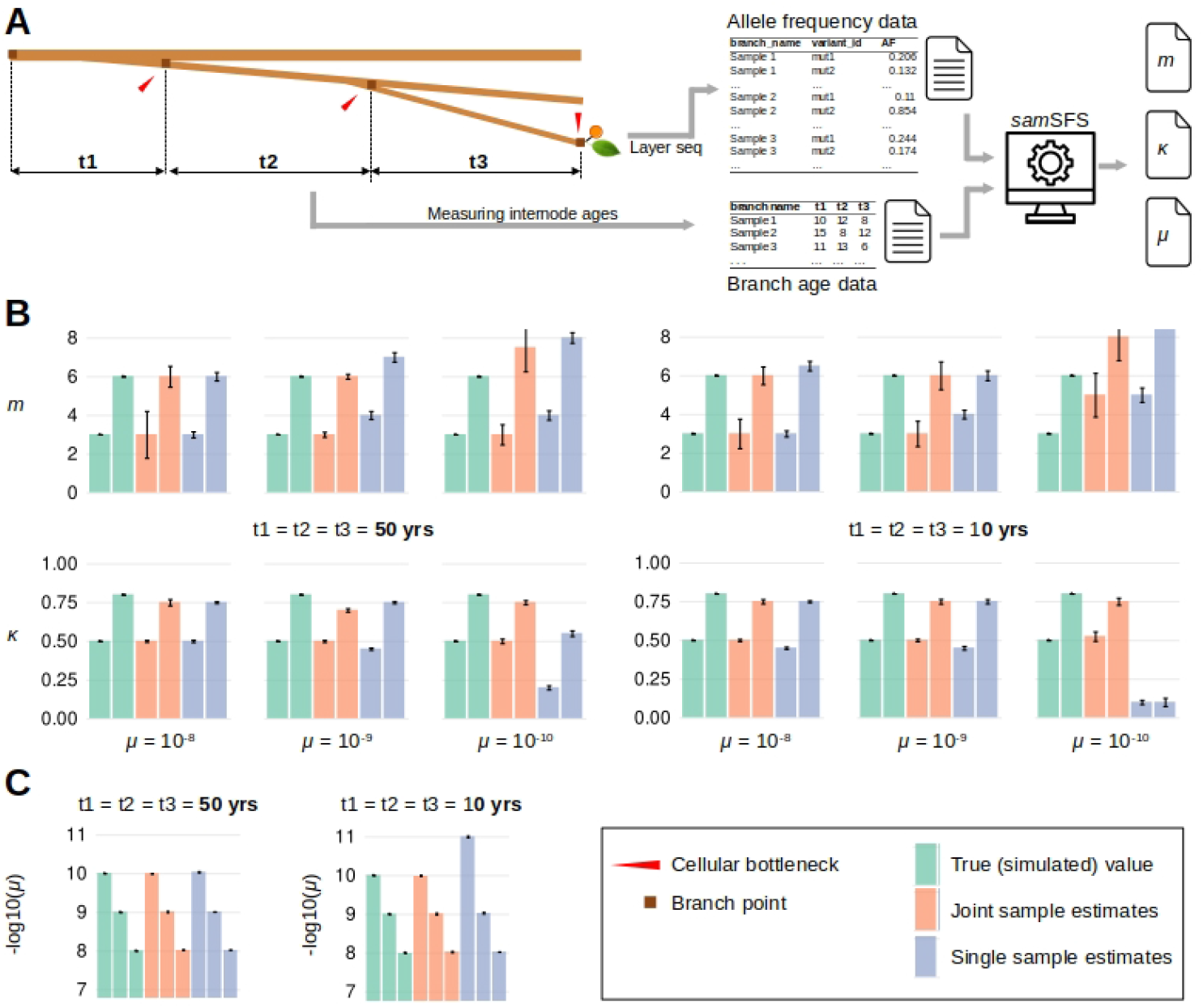
Overview *sam*SFS software and simulation study. **A**. Inputs/outputs of the *sam*SFS software (https://github.com/jlab-code/samSFS). The main input are two files: 1. Branch age data consisting of internode age measurements, and 2. Allele frequency data obtained from sequencing of layer-enriched fruit or leaves, or from direct sequencing of SAM cell layers, if the later becomes technically feasible. The output are statistical estimates of the parameters *m, κ* and *µ*. **B**. Simulations were conducted using the branching topology shown in A., which consists of three nested branching events and internode ages *t*_1_, *t*_2_ and *t*_3_. All results are presented as means of replicate simulation runs (+/- SE), assuming random AM placement and a genome size of *G* = 240 · 10^6^ Mb. The full simulation results can be found in Tables S2 and S3. Shown are parameter estimates of *m* and *κ* for three different simulated mutations rates *µ* = 10^−8^, *µ* = 10^−9^ and *µ* = 10^−10^, as well as for two different branch internode ages *t*_1_, *t*_2_, *t*_3_ = 50 (left panel) and *t*_1_, *t*_2_, *t*_3_ = 10 (right panel). In each barplot, single sample estimates (orange) and joint sample analysis (blue) are compared with the simulated (true) values of the parameters (green). **C**. Estimates of the mutation rate parameter *µ* for the two different branch internode ages *t*_1_, *t*_2_, *t*_3_ = 50 (left panel) and *t*_1_, *t*_2_, *t*_3_ = 10 (right panel).

Each simulation scenario was defined by a combination of ground truth parameter values, a set of internode time intervals **t** = (*t*_1_, *t*_2_, …, *t*_*n*_), and a binning resolution *u* over the allele frequency range. We considered a branching scenario with three nested events, where the internode intervals were either 10 or 50 years, corresponding to total tree ages of 30 or 150 years, respectively. Mutation accumulation along branches was determined by three different mutations rates *µ* ∈ {10^−8^, 10^−9^, 10^−10^} per haploid genome per year. We assessed inference performance across combinations of *κ* ∈ {0.5, 0.8}, *m* ∈ {3, 6}, and a range of mutation rates including biologically realistic values. For each parameter combination, we generated 100 independent replicate datasets and applied the inference procedure to each. The full numerical results of the simulation can be found in TableS2 and TableS3.

We found that in the single-sample analysis, parameter estimates were highly accurate in most parameter regimes. However, the estimates became inaccurate when the mutation rate was as low as *µ* = 10^−10^, a problem that was further enhanced when branch internode times were short i.e. *t*_1_, *t*_2_, *t*_3_ = 10. Clearly, the reason for this is that the low mutation numbers in these situations hold little information about the shape and scale of the *sam*SFS. As explained in section: Modelling the *sam*SFS), this issue can be partly corrected with the joint inference approaches, where multiple samples are batched in a weighted interference approach. This improvement can be seen, for example, when looking at parameter *κ* when *µ* = 10^−10^ and *t*_1_, *t*_2_, *t*_3_ = 10. Further improvements are possible when aggregating the internode and mutation count data of multiple samples and fitting an aggregate likelihood. That is, if we aggregated five samples from the *t*_1_, *t*_2_, *t*_3_ = 10 regime, the data would become equivalent to a *t*_1_, *t*_2_, *t*_3_ = 50 regime. The information boost obtained with this strategy is illustrated when comparing the estimates of *κ* or *µ* in (Fig. 4). Given these analysis options and the fact that the per year somatic mutation rates reported in long-lived plant species are typically in the order of 10^−9^ (Johannes (2024)), we were confident that our method could be successfully applied to real data.

## Data application: Apricot tree

We applied the *sam*SFS software to recent data from the study of Goel et al. (2024)). The authors analyzed somatic mutations in multiple branches of an apricot tree (*Prunus armeniaca cv. Rojo Pasión*) (Fig.5A). In contrast to previous tree studies (Duan et al. (2022), Hanlon et al. (2019), Hofmeister et al. (2020), Wang et al. (2019), Orr et al. (2020), Perez-Roman et al. (2022), Schmid-Siegert et al. (2017), Xie et al. (2016), Satake et al. (2023)), the they performed DNA sequencing of two distinct tissues that are known to be derived from different SAM cell layers (fruit peel: L1-derived; mesocarp: L2-derived). Their analysis revealed that *de novo* meristematic mutations appear in a layer-specific fashion and that L1 layer mutations are much more frequent than L2-derived mutations, though no mutation rates were reported.

**Figure 5:**
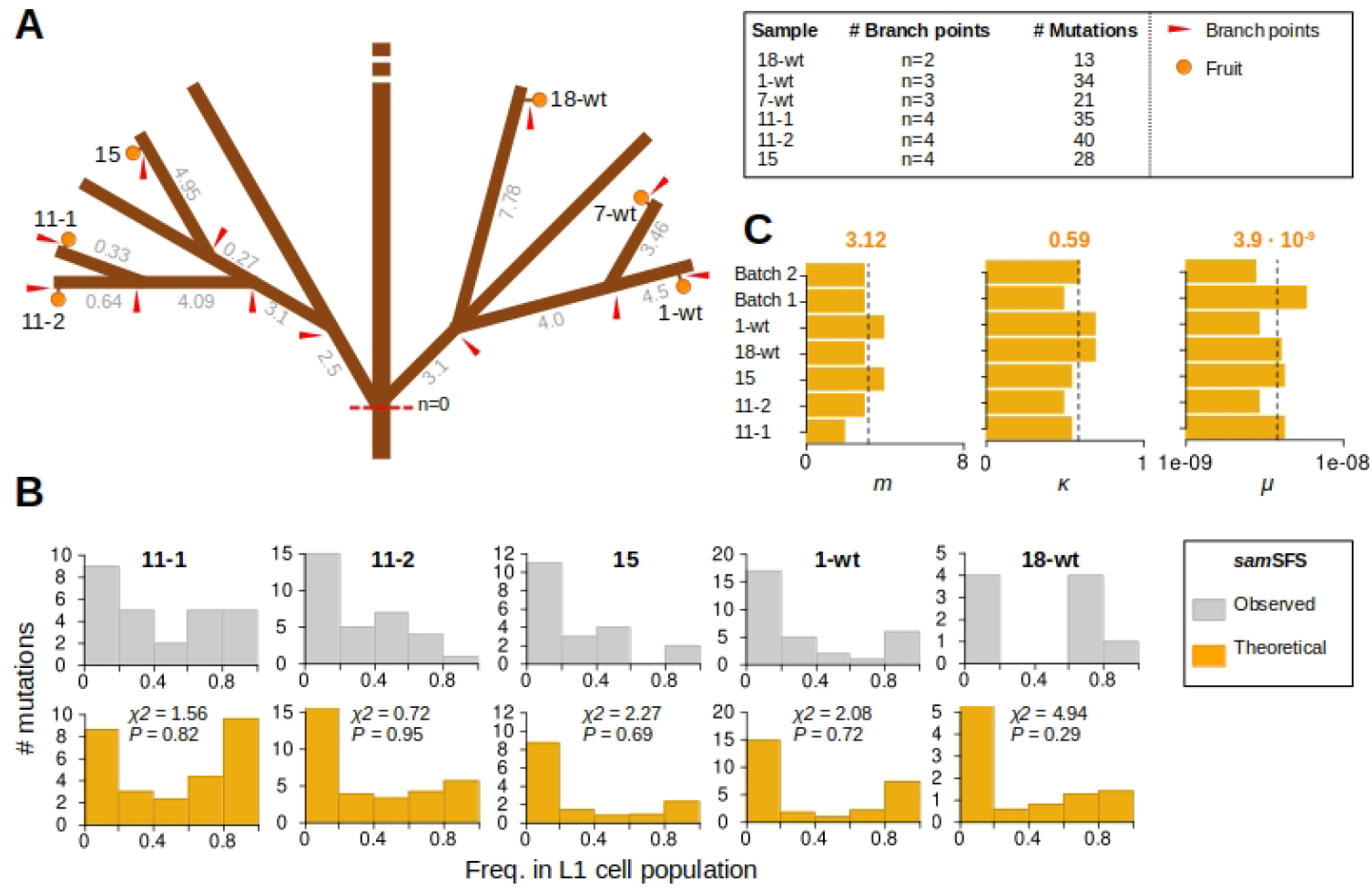
Data analysis of apricot tree: **A**. Topology of the apricot tree (*Prunus armeniaca cv. Rojo Pasión*) analyzed by Goel et al. (2024). Layer-enriched DNA sequencing was performed for six different branches (11-2, 11-1, 15, 1-wt, 7-wt, 18-wt). Branching bottlenecks are indicated by red arrows and were counted from the tree base (*n* = 0) to the location of each sample on the terminal branches. Gray numbers denote the approximate age (in years) for all internode segments. **B**. Top panel: Observed *sam*SFS; Bottom panel: estimated theoretical *sam*SFS. The goodness-of-fit statistics and their corresponding *P* values are shown for this model. **C**. Estimates of the concentration parameter (*κ*), the size of the apical stem cell population (*m*) and the SAM L1-layer haploid mutation rate per bp per year (*µ*) for each sample. Note that sample 7-wt was an outlier for estimates of *m* (see Table S4), and was therefore omitted here from the single sample plot; however, we did include that sample in the joint analysis (i.e. see Batch 2).

Building on these data, we focused on variant calls of L1-derived mutations from six branches (Fig.5A, Methods). The observed SFS of each of these branches is displayed in Fig.5B (Method). We performed both the single and joint sample analysis. For the joint analysis we created two batches, where the first included samples 11-2, 11-1, 15 and the second 1-wt, 7-wt, 18-wt. Overall, both analysis approaches revealed very similar results. On average, we estimated that ASC population size is (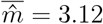 (*SE* = 0.26, a number that is in line with live imaging observations in Arabidopsis and tomato (3-4 ASC cells) Burian et al. (2016)). Moreover, the model estimated a mutation rate of 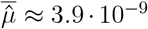 (*SE* = 4.2 · 10^−10^ per bp per year. This value is in the range of previous estimates in the Prunus family (e.g. *Prunus persica*: *µ* = 0.82 · 10^−9^; *Prunus mume*: *µ* = 2.2 · 10^−9^) (Wang et al. (2019)). Finally, our estimate for the concentration parameter was 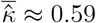 (*SE* = 0.032). This value is somewhat low, and suggests precursor sampling is not entirely restricted to a single SAM clonal sector, on average. However, a technical caveat with the estimates of *κ* is that both high- and low-frequency variants are likely undersampled in the post-processed sequencing data. Under-sampling of high-frequency variants could stem from layer contamination during tissue dissection. Under-sampling of low frequency mutations, on the other hand, may be due to stringent filtering of rare variant calls that resemble sequencing noise. The net effect is that the concentration parameter is biased downward, so that it mimics a relaxed drift scenario. It should be possible to correct these issues in future empirical studies, for example by using layer-specific marker genes in combination with cell sorting prior to sequencing, and by fine-tuning pre-processing thresholds for variant calling.

Despite these caveats, our model agrees well with the observed data (see goodness-of-fit test in Fig.5B), and the parameter estimates for *µ* and *m* are consistent with those reported in other species. Hence, a statistical analysis of the *sam*SFS in combination with DNA sequencing data appears to be feasible methodological approach, which can yield estimates of meristematic parameters that are difficult to obtain with other methods.

## Conclusion and outlook

Here we developed a theoretical model that describes how the frequency of *de novo* mutations in apical stem cells (ASCs) evolve in the SAM cell population due to neutral somatic drift during repeated branching. The process is a function of the ASC population size (*m*), the strength of the cellular bottleneck during branch initiation (*κ*), and the stem cell mutation rate (*µ*). Using the model, we derived a method for computing the mutation site frequency spectrum of the SAM (*sam*SFS), and showed that it can be applied to layer-enriched DNA sequencing data for parameter estimation. This is a useful result because there are currently no alternative methods for measuring these parameters *in vivo*, and certainly not simultaneously.

Perhaps the most important outcome of our analysis is that the model aligns closely with the observed *sam*SFS in our proof-of-principle data application. This strong agreement suggests that the basic neutral mutation-drift framework is sufficient to explain the observed site frequency spectrum, without requiring more complex biological mechanisms. Notably, this challenges the necessity of invoking additional processes such as layer exchange or cell lineage selection—mechanisms that have featured prominently in several theoretical studies (e.g., Iwasa et al. (2023), Klekowski et al. (1985), Pineda-Krch and Lehtila (2002)) but may play a limited role in shaping within-SAM genetic diversity in practice.

Although our model was fit to data on somatic mutations, it is readily adaptable to other molecular markers—particularly spontaneous somatic epimutations. These are mitotically heritable changes in CG methylation that occur stochastically during cell divisions (Johannes and Schmitz (2019)). Importantly, their rate of accumulation is several orders of magnitude higher than that of somatic genetic mutations (van der Graaf et al. (2015), Hazarika et al. (2022), Denkena et al. (2021), Schmitz et al. (2011), Becker et al. (2011)). As a result, they offer a denser signal for analysis and could significantly enhance statistical power for model inference. Indeed, recent work has shown their potential to provide fine-scale insights into lineage dynamics over time (Yao et al. (2021), Yao et al. (2023)).

However, an important caveat concerns the use of bulk tissue samples. Given the theoretical basis of our model, we suspect that bulk sequencing may obscure accurate inference, as the resulting *sam*SFS becomes a composite of layer-specific variant allele frequencies (VAFs), with unknown and potentially variable mixing proportions. This mixing introduces ambiguity that is difficult—if not impossible—to deconvolute without additional information. For this reason, layer-enriched sequencing is essential for reliable parameter estimation, as emphasized earlier and supported by recent findings (e.g., Goel et al. (2024)).

While our data application was restricted to apricot, the method has clear applications beyond trees. It can be applied, generally, to any plant species that uses branching as a developmental or reproductive strategy. This also includes species that propagate clonally via ramets, suckers, tillers, etc. Examples include clonal populations of the seagrass (*Zostera marina*) (Yu et al. (2019)), or the large Aspen (*Populus tremuloides*) clones of the Pando forest (Pineau et al. (2024)). Since our model assumes a neutral mutation-drift processes, it could serve as a starting point to detect evidence for cell lineage, module or ramet selection in natural settings. Possible model extensions could include the direct incorporation of a selection parameter, and the possibility of L1/L2/L3 stem cell layer-exchange, as these processes may be more important over long time-scales of clonal evolution.

A key experimental challenge in applying our method is the need to know branch or internode ages, which are typically obtained through labor-intensive techniques like coring—especially burdensome when applied to multiple branches. A promising alternative, though requiring further validation, is to use branch or internode length as a proxy for age. This may be particularly useful when the main goal is estimating meristematic parameters such as *κ* and *m*. For this proxy to be effective, mutation accumulation per unit length must scale similarly to that per unit time. Evidence supporting this approach comes from the apricot dataset, where only total tree age was available, yet internode ages were inferred from measured lengths—yielding mutation rates consistent with those in the literature.

We expect that further tests of our model on a broader range of plant species and data modalities will lead to further refinements. Such efforts promise to accelerate our basic understanding of the somatic evolution of stem cell mutations in long-lived plants.

## Materials and Methods

### Implementation of the wrapped Cauchy distribution

To implement the theoretical models described in the main text, we needed to assume a specific form for the underlying wrapped distribution, *f* (*θ*; *ν, κ*). A flexible choice for modeling precursor sampling is the wrapped Cauchy distribution. Its probability density function (PDF) is given by:

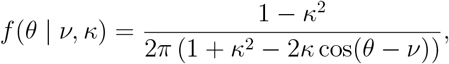

where *κ* ∈ [0, 1). The bounds on *κ* facilitate an easy interpretation of the strength of the bottleneck during branching.

### Exploratory analysis of fixation times

To explore fixation times, we restricted the analysis to a realistic parameter range of values of *κ* such that 90% of the cumulative probability density of the wrapped distribution is contained within one-half of the unit circle. This constraint reflects microscopy studies showing that AM initiation and precursor sampling are localized phenomena, occurring in “narrow” regions along the SAM periphery. To determine the minimum *κ* value that meets this condition, we computed the cumulative probability over the interval [−*π/*2, *π/*2] as a function of *κ*. Specifically, the cumulative probability for a given *κ* was calculated by integrating the probability density function (PDF) of the wrapped Cauchy distribution around the mode *ν*:

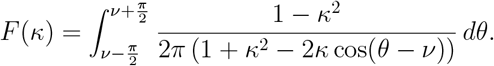

This computation was repeated over a grid of increasing *κ* values until the condition *F* (*κ*) ≥ 0.90 was met. Based on the wrapped Cauchy distribution, the minimum *κ* value that satisfies this requirement is approximately 0.7. We used a similar approach to generate Fig.2B. There, the goal was to find the minimum value of *κ* corresponding to different *m*, so that the above probability criterion was met in each case. Hence, for a given *m*, integration was over the interval [*ν* − *π/m, ν* + *π/m*].

### Implementation of the likelihood-based model inference

We estimated the parameters Ψ = (*m, κ, µ*) by maximizing the log-likelihood function:

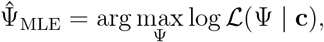

where

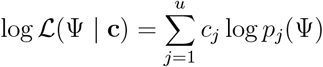

is defined by the multinomial model for the binned variant allele frequencies (VAFs). We performed the maximization over a structured, discrete grid of candidate values for Ψ. This grid-based approach was chosen for three reasons. First, the typical number of mutations per sample is modest, which can yield a rough or multimodal likelihood surface. Second, the parameter space is low-dimensional and bounded: *m* is discrete, *κ* ∈ (0, 1), and *µ* spans a narrow, log-scaled range. Third, an exhaustive evaluation over the grid ensures robust estimation and facilitates downstream model comparisons.

The inference procedure was implemented in `R`, with parallelization across samples and embedded `C++` routines for forward-in-time simulation of the multinomial mixture density:

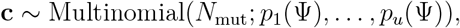

for each candidate Ψ. For joint analysis across multiple samples, we used a weighted average of the sample-specific log-likelihoods, where weights *α*_*s*_ reflected the number of observed mutations in each sample.

To characterize uncertainty and parameter identifiability, we used the top *Q* highest-scoring grid points Ψ_*i*_ = (*m*_*i*_, *κ*_*i*_, *µ*_*i*_) to summarize the structure of the likelihood surface. For each parameter combination Ψ_*i*_, we computed the corresponding log-likelihood:

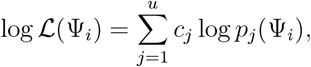

or, in the case of joint estimation, a weighted sum across samples. We then assigned normalized weights to each grid point:

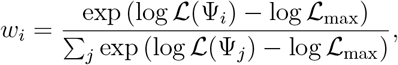

where log ℒ_max_ was the maximum log-likelihood among the top *Q* grid points. These weights reflected the local structure of the likelihood surface, analogous to a softmax transformation, and were used to compute likelihood-weighted summaries of each parameter (e.g., for *m*):

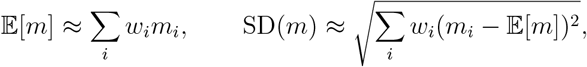

and similarly for *κ* and *µ*. Weighted medians were defined as the first value at which the cumulative sum of weights exceeded 0.5. We also computed the average log-likelihood across the top *Q* grid points:

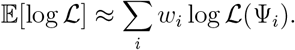

This approach provided a smooth, likelihood-informed characterization of parameter values that was robust to flat or multimodal regions in the parameter space. While our implementation is purely likelihood-based, this framework can be naturally extended to a Bayesian setting by introducing prior distributions over *m, κ*, and *µ*, and performing inference using, for example, Markov Chain Monte Carlo (MCMC) sampling over the posterior.

### Simulation study

The implemented a simulator to assess whether our inference framework could recover the true parameters under different parameter regimes. The simulator is written R with extensive parallelization and an embedded C++ routine, and can be found at https://github.com/jlab-code/samSFS. User-friendly demo code is provided for testing.

### *sam*SFS analysis in apricot

We reanalyzed data from the study of Goel et al. (2024), who sequenced fruits from six different branches (11-1, 11-2, 15, 7-wt, 1-wt, 18-wt) of an apricot tree. A unique aspect of this study is that, prior to sequencing, the authors dissected apricot fruits into two distinct tissues that are known to be derived from different SAM cell layers (fruit peel: L1-derived; mesocarp: L2-derived). For each branch, we obtained a dataset of *de novo* mutations (Table S2). To guard against variant over-filtering, we worked with a rawer version of the final dataset the authors used in their original publication. To obtain this data, the following filtering steps were applied: Reads were trimmed, aligned, and deduplicated. Positions with at least three reads supporting the variant allele and displaying mapping quality ≥10 and base quality ≥30 were retained. Further positions were removed if they: were heterozygous between the two haplotypes of the assembly, had too low/high read depth in either L1 or L2 sample, or did not have sufficient variant allele frequency (VAF) variation between the layers, represented positions that were mutated in all of the leaf samples. Since apricot is diploid the VAF values of each mutation were multiplied by two to obtain a proxy of the proportion of cells carrying the mutation in a given bulk sample. Adjusted VAF values *>*1 were set to 1.

To construct the observed *sam*SFS, we divided the vector of VAFs into *u* = 5 equally-sized intervals over the range [0, 1] and counted the values in each interval. The interval boundaries [*d*_1_, *d*_2_, …, *d*_*u*+1_] were defined, with *d*_1_ set at 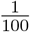. For each interval [*d*_*q*_, *d*_*q*+1_), we counted the VAFs, *v*_*r*_, that satisfied *d*_*q*_ ≤ *v*_*r*_ *< d*_*q*+1_. These counts formed a vector, representing the observed SFS (Fig.5B). To be able to fit the theoretical *sam*SFS to each of the samples, we needed information about the parameter *n* and as well as the internode time intervals *t*_1_, …, *t*_*n*_. Starting from the observed tree topology, we determined *n* for each sample, by counting the number of bottlenecks along a branching path from the tree base to the location of that sample. The tree used in this study was 11 years old. However, the ages of the internode segments was not available, but only provided in terms of “meters of growth”. We therefore converted the internode segments to approximate time intervals *t*_1_, …, *t*_*n*_ (in years). To do this, we assumed a constant growth rate and expressed the meter-length of each segment as a proportion of the total meter-length of the branching path from the tree base to the sampled fruits. These proportions were then multiplied by the total tree age to obtain the approximate ages of each segment in terms of years.

## Acknowledgments

I thank Korbinian Schneeberger and Manish Goel for sharing and explaining their semi-filtered datasets, and Agata Burian for her quick feedback on a draft version of this manuscript.

